# Validation of helical symmetry parameters in EMDB

**DOI:** 10.1101/2025.05.18.654508

**Authors:** Daoyi Li, María Muñoz Pérez, Xiaoqi Zhang, Jiaqing Li, Wen Jiang

**Affiliations:** Department of Biological Sciences, Purdue University, West Lafayette, IN 47907; Department of Biochemistry and Molecular Biology, The Pennsylvania State University, University Park, PA 16802; Center for Structural Biology, Huck Institutes of the Life Sciences, The Pennsylvania State University, University Park, PA 16802

**Keywords:** cryo-EM, validation, helical symmetry, Electron Microscopy Data Bank

## Abstract

Helical symmetry is a structural feature of many biological assemblies, including cytoskeletons, viruses, and pathological amyloid fibrils. One unique metadata for helical structures is the helical parameters, twist and rise. With the increasing number of helical structures being resolved through cryo-EM and deposited in the EMDB, there is a growing possibility of errors in the metadata associated with these entries. During our cryo-EM analysis of protein amyloids and the development of helical analysis tools, we realized that many deposited helical parameters appear inconsistent with the associated density maps. Herein, we have developed a comprehensive validation process that examines the consistency of these parameters by combining high-throughput computational evaluation with manual verification. Multiple errors were identified and corrected for ∼14% of the total entries, including missing parameters, swapped twist and rise values, incorrect sign of twist angles, partial symmetries, and bona fide errors. Our validation code, workflow, and the validated parameters are publicly available.

**Synopsis:** This article reports a systematic validation of the consistency between the deposited helical parameters and the density maps of helical structures in EMDB. Multiple parameter errors were identified and corrected.

## 1. Introduction

Helical symmetry is a structural feature of many biological assemblies, including cytoskeletons (Chakraborty *et al*., 2020), viruses (González *et al*., 2021), and pathological amyloid fibrils (Fernandez *et al*., 2024; Hallinan *et al*., 2021). Determining helical structures at atomic level is crucial for understanding their biological functions and can guide drug discoveries. Cryogenic electron microscopy (cryo-EM) is a powerful tool for resolving structures at near-atomic resolutions. Since the first helical structure was solved using EM in 1968 (De Rosier & Klug, 1968), EM has been widely employed in the structural study of helical assemblies.

Similar to point group symmetries, structures with helical symmetry are comprised of a single asymmetric unit that replicates according to the symmetry order (Tykač *et al*., 2021). In contrast to point group symmetries where the symmetry transformations are pure rotations around a fixed point or axis, helical symmetries involve both rotational and translational transformations, specifically the rise (translation along the helical axis), twist (rotation around the helical axis), and axial symmetry (Diaz *et al*., 2010) (Fig. 1A). Instead of the discrete rotational orders, such as two-fold, three-fold, *etc*., for the point group symmetries, helical rise and twist can take arbitrary values. Helical symmetry is thus a family of symmetries with infinite options, and helical indexing, *i*.*e*., accurately identifying the rise and twist values, is a challenging but essential task for *de novo* determination of helical structures.

**Figure 1.**
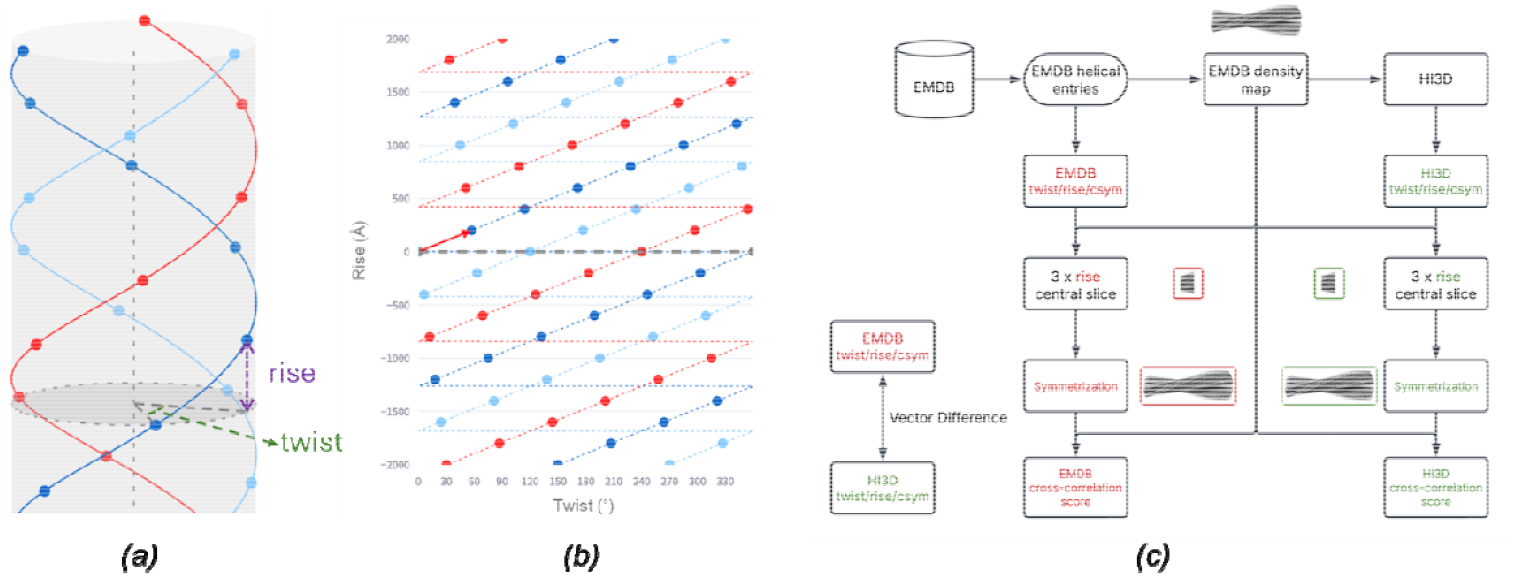
Workflow for the validation of helical parameters in EMDB. *(a)* A diagram illustrating helical symmetry. *(b)* The 2D lattice resulted from unwrapping the helical lattice shown in *(a)*. The lattices shown in *(a)* and *(b)* were generated using the HelicalLattice Web app with twist=57°, rise=200 Å, C3 axial symmetry, and radius=500 Å. To reproduce these plots, visit https://helical-lattice.streamlit.app/?diameter=1000&length=4000&rise=200&twist=57&csym=3&rot=10&tilt=10&draw_cylinder=1&draw_axis=1&marker_size=10. *(c)* Validation workflow. This workflow illustrates the validation process for the helical parameters (twist, rise, and axial symmetry) of the helical structure entries in EMDB. The deposited helical parameters (red) are retrieved from the metadata of EMDB entries, while the validated helical parameters (green) are determined using HI3D. The vector difference and relative difference between these two sets of helical parameters are compared. To assess the validity of each parameter set, the corresponding density maps were central-sliced at three times the helical rise and were symmetrized into full-length helical densities based on either the deposited parameters or the HI3D parameters. The similarity between the symmetrized map and the original density map was evaluated using a cross-correlation scoring metric. The parameter set yielding a significantly higher cross-correlation score is considered a better representation of the density map.

With an increasing number of EM structures deposited in the Electron Microscopy Data Bank (EMDB), there is a growing possibility of errors in the deposited structures and the associated metadata (The wwPDB Consortium, 2024; Chiu *et al*., 2021). The helical assemblies account for ∼4.5% of the entries in the EMDB (EMDB, 2025). During our cryo-EM analysis of protein amyloids and the development of helical analysis tools (Sun *et al*., 2022; Li *et al*., 2025; Li & Jiang, 2023), we realized that many deposited helical parameters appear inconsistent with the associated density maps. As current validation pipelines predominantly focus on single-particle reconstructions, there is a lack of robust validation of the specialized parameters associated with helical symmetry (Wang *et al*., 2022). A systematic approach, combining high-throughput computational evaluation with manual verification, will improve the reliability of the helical parameters associated with EMDB density maps, enabling more accurate biological interpretations, preventing further propagation of the errors, and benefiting downstream structural analysis and machine learning using the helical structures and parameters.

The helical symmetry parameters are traditionally determined by indexing the layer line patterns in the power spectra of 2D images before 3D reconstruction (Diaz *et al*., 2010). When a 3D density map is available, we previously reported a real-space indexing method, HI3D (Sun *et al*., 2022), which converts a 3D helical structure into a 2D lattice via cylindrical projection and recasts helical indexing as a unit cell determination task for a 2D lattice (Fig. 1A,B). While the primary goal of HI3D was to allow *de novo* helical indexing using a single particle or tomographic 3D reconstruction with no helical symmetry being imposed, the capability of automatically estimating the helical symmetry from the 3D density map makes HI3D a convenient tool to evaluate if the deposited helical parameters are consistent with the corresponding density maps in EMDB.

In this study, we present a comprehensive pipeline designed to validate the helical parameters of all helical structure entries in the EMDB (Fig. 1C). We first used HI3D to systematically analyse each helical entry and automatically detect the helical parameters. Subsequently, we compared the deposited helical parameters with those identified by HI3D. Entries that could not be reliably validated through this automated process were manually examined to ensure accuracy.

Our analyses revealed multiple issues with helical parameters deposited in the EMDB, including no values, swapped twist/rise values, incorrect twist sign, partial symmetry, and incorrect twist/rise values. For the entries using partial symmetries, varying degrees of improvements in resolution were obtained when the maps were symmetrized using the full symmetry identified by our validation.

## 2. Methods

### 2.1. EMDB entries with helical symmetry

The EMDB includes structures determined with different methods. In this work, we focused exclusively on density maps solved using helical reconstruction, where the *structure determination method* in the metadata is *helical*. The list of helical structures, the density maps, and the deposited helical parameters, including rise, twist, and axial symmetry, were programmatically retrieved from EMDB. There are 2,025 helical structures in EMDB using a cutoff date of April 25, 2025.

### 2.2. Automated helical indexing with HI3D

As depicted in Fig. 1C, we have developed an automated pipeline to systematically validate the helical parameters of all helical structures in EMDB utilizing HI3D. Since HI3D was originally developed as an interactive Web app, we adapted the core computational routines into a batch script. For non-amyloid helical structures, we used the default HI3D parameters, with an angular step of 1° and an axial step of 1 Å. However, for amyloid structures, where the twist angle and rise are relatively small, we used finer samplings with an angular step of 0.5° and an axial step of 0.2 Å to improve accuracy.

### 2.3. Comparing the deposited helical parameter and the helical parameter determined by HI3D

After obtaining the helical parameters using HI3D, we systematically compared the deposited helical parameters with those determined by HI3D. We introduced three distinct metrics to evaluate the differences between the two sets of helical parameters.

The first metric computes the normalized differences. The differences between the original deposited helical parameters (*θ*_*ori*_, *z*_*ori*_) and those determined by HI3D (*θ*_*HI3D*_, *z*_*HI3D*_) can be calculated as:

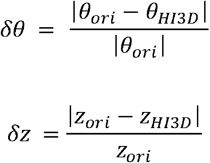

Where *δθ* and *δz* represent the normalized differences in twist and rise, respectively. However, this method treats twist and rise separately without considering their correlation and distinct nature (e.g. rotation vs shift). Additionally, it does not account for the diameter of the helical structure, making it less applicable for general comparisons across different structures.

The second metric evaluates differences by incorporating the twist, rise, and radius of the helical structure. To estimate the radius (*r*), we calculated the radial profile of the densities around the helical axis. The helical parameters were then converted into vectors as follows:

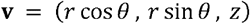

The difference between the two sets of helical parameters was then calculated as the Euclidean distance between their respective vectors of helical parameters:

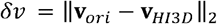

If this vector difference is smaller than the reported resolution, the two parameter sets are considered similar. However, this approach is insensitive to cases where the original helical parameters are small but differ in sign. For instance, an amyloid structure with a deposited twist value of 0.4° and a HI3D-detected value of -0.4° would indicate different handedness, yet the vector difference would remain small.

The final metric involves symmetrizing the original deposited density map using both sets of helical parameters. The transformation ***T***_***i***_ is defined as:

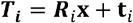

The transformation of the asymmetric unit is the combination of the rotational group ***R***_***i***_ and translational **t**_***i***_ along the helical axis. Through this transformation, the symmetrized density could be defined as:

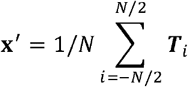

where **x**′ is the final symmetrized density map and *N* is the number of symmetry transformations. We then compute the cross-correlation scores between the original density map and the symmetrized density maps generated using either the original deposited helical parameters (*CC*_*ori*_) or the HI3D helical parameters (*CC*_*HI3D*_). If the cross-correlation score with one set of helical parameters is significantly higher, it indicates that this parameter better explains the structure.

These three metrics provide a comprehensive evaluation of the helical parameters, ensuring consistency between the deposited density map and associated metadata.

### 2.4. Manual validation

In some cases, the cross-correlation score between the symmetrized map and the original map, using both sets of helical parameters, can be low (< 0.5), and the two sets of helical parameters differ significantly. In such instances, manual validation of the helical parameters is necessary.

There are multiple possible reasons for this discrepancy. One possibility is that the deposited helical parameters are incorrect, and simultaneously, HI3D fails to detect the correct helical parameters using the default settings. This issue can often be resolved by testing multiple combinations of angular step and axial step parameters in HI3D. Another scenario occurs when the deposited map in EMDB only includes a partial density map, for example, an asymmetric unit of the helical reconstruction, or a focused reconstructed map. In such cases, there is not sufficient information for HI3D to estimate the helical parameters accurately, as it relies on the correlation of multiple asymmetric units for reliable estimation. We have noted these entries as not validated in our final table of validated helical parameters.

Another complication is that the order of the voxels in the map file of 66 entries does not follow the standard axes order (fastest varying x, y, slowest varying z). Our code checks the axes order and automatically reorders the voxels to the standard axes order. However, some entries could not be automatically corrected as their voxel order is inconsistent with the axis order specified in the MRC/CCP4 map header. Thus, we tried other combinations of axis orders to find one that would orient the helical axis along the Z axis and subsequently imported the corrected map into HI3D.

### 2.5. Partial symmetry and resolution evaluations using the symmetrized half maps

Another scenario arises when the cross-correlation scores for both the deposited and HI3D-derived helical parameters are similar, but the relative and vector differences are large. This situation suggests that both sets of helical parameters are correct, but one set might have only used partial helical symmetry.

For example, for a helical structure with a twist of θ and a rise of *z*, a reconstruction using a twist of n*θ and a rise of n\**z* (n=2, 3, …) still yields a correct structure. However, such a reconstruction does not necessarily achieve the best structural quality, as it only utilizes a fraction of the full symmetry that results in less coherent averaging. To assess whether this is the case, we evaluate if the two sets of helical parameters are related by an integer multiplier and visually examine if both helical parameter sets point to lattice points in the HI3D lattice plot. Furthermore, we symmetrize the two deposited half-maps using both the deposited and the validated helical parameters and evaluate whether the resolution improves according to the Fourier shell correlation of symmetrized half-maps.

### 2.6. Structural analysis and visualization

The helical and 2D lattices shown in Fig. 1A,B were generated using the HelicalLattice Web app (https://jianglab.science.psu.edu/helicallattice). The 3D maps were visualized using ChimeraX (Pettersen *et al*., 2021). For optimal visualization, some maps were auto-sharpened using Phenix (Terwilliger *et al*., 2018). JSPR was used to match structure factor profiles across different maps (Sun *et al*., 2020).

### 2.7. Implementation and code/result availability

The validation scripts were implemented in Python, utilizing the HI3D algorithm as the core method. The source code and the validation results are freely available on GitHub (https://github.com/jianglab/EMDB_helical_parameter_validation).

## 3. Results

### 3.1. Differences between deposited and validated helical parameters

The complete list of validated helical symmetry parameters for all helical structures in EMDB is available in Appendix A. We compared the overall differences between the deposited and validated helical parameters. The histogram shown in Fig. 2A illustrates the distribution of vector differences (in Å) between the deposited helical and the newly validated values. While most entries exhibit relatively minor differences, a noticeable tail extends to large differences. This indicates that a considerable number of discrepancies exist between the deposited and validated helical parameters.

**Figure 2.**
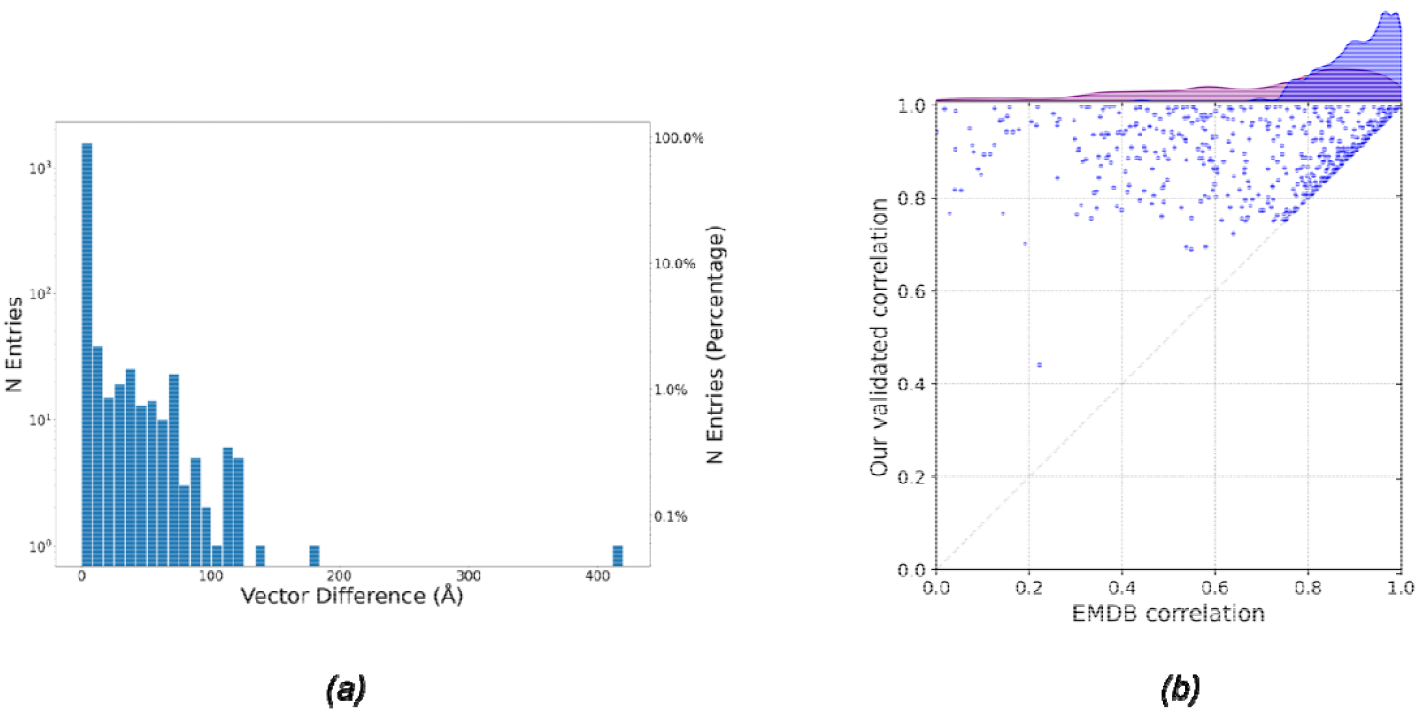
*(a)* A histogram displaying the distribution of vector differences between the deposited and scatter plot comparing the cross-correlation scores between the symmetrized map and the original density map, using either the originally deposited helical parameters (X-axis) or the validated helical parameters (Y-axis). Only entries with improvement of the cross-correlation score between the two sets of parameters were included in the plot. The histograms of the correlations from the deposited helical parameters (purple) and the validated helical parameters (blue) were displayed on top of the plot. The histogram from the validated helical parameters was noticeably skewed to the right (i.e. better correlations).

Fig. 2B compares the cross-correlation scores between symmetrized and original maps for both parameter sets, where only the entries with discrepant helical parameters are included in the plot. Each point represents a single EMDB entry, with the x-axis showing the cross-correlation score when using the deposited parameters and the y-axis showing the cross-correlation score when using the validated parameters. Points above the diagonal line indicate cases where the validated parameters produce a higher cross-correlation score than the originally deposited parameters. The overall trend reveals that the validated parameters improve or at least maintain the correlation score. This result suggests that the validation process can yield the helical parameter that is more consistent with the deposited density map.

### 3.2. Typical errors in the deposited helical parameters

A systematic analysis of helical structure entries in the EMDB revealed that a considerable number of entries contain errors in their deposited helical parameters. These errors can potentially mislead structural interpretation and affect downstream analyses. The types of typical errors and examples of these errors are summarized in Table 1, and their corresponding visual representations using HI3D lattice plots are shown in Fig. 3.

**Table 1.**
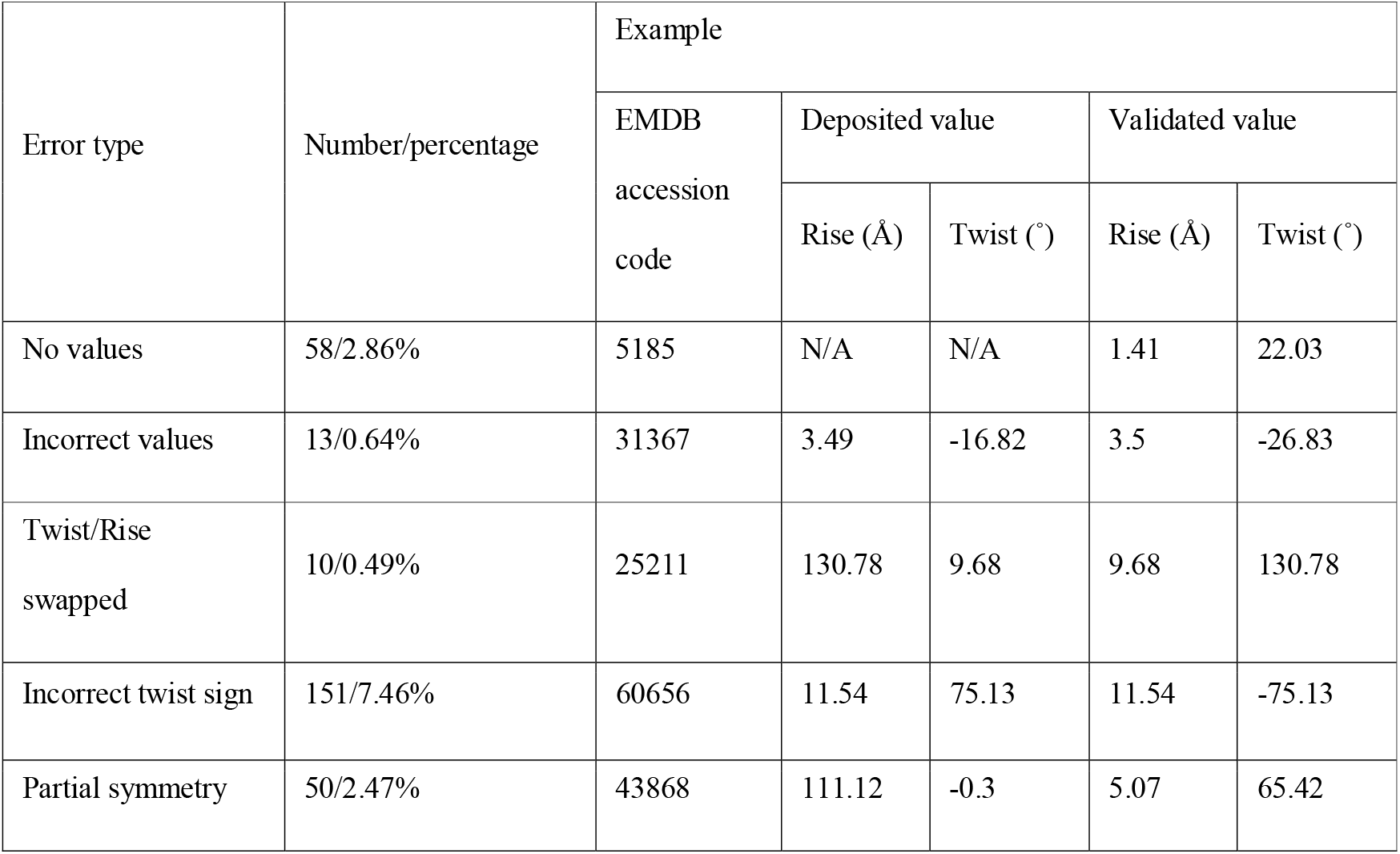
Types of errors in the deposited helical parameters in EMDB.

| Error type | Number/percentage | Example |  |  |  |  |
| --- | --- | --- | --- | --- | --- | --- |
|  |  | EMDB<br>accession<br>code | Deposited value |  | Validated value |  |
|  |  |  | Rise (Å) | Twist (°) | Rise (Å) | Twist (°) |
| No values | 58/2.86% | 5185 | N/A | N/A | 1.41 | 22.03 |
| Incorrect values | 13/0.64% | 31367 | 3.49 | -16.82 | 3.5 | -26.83 |
| Twist/Rise swapped | 10/0.49% | 25211 | 130.78 | 9.68 | 9.68 | 130.78 |
| Incorrect twist sign | 151/7.46% | 60656 | 11.54 | 75.13 | 11.54 | -75.13 |
| Partial symmetry | 50/2.47% | 43868 | 111.12 | -0.3 | 5.07 | 65.42 |

**Figure 3.**
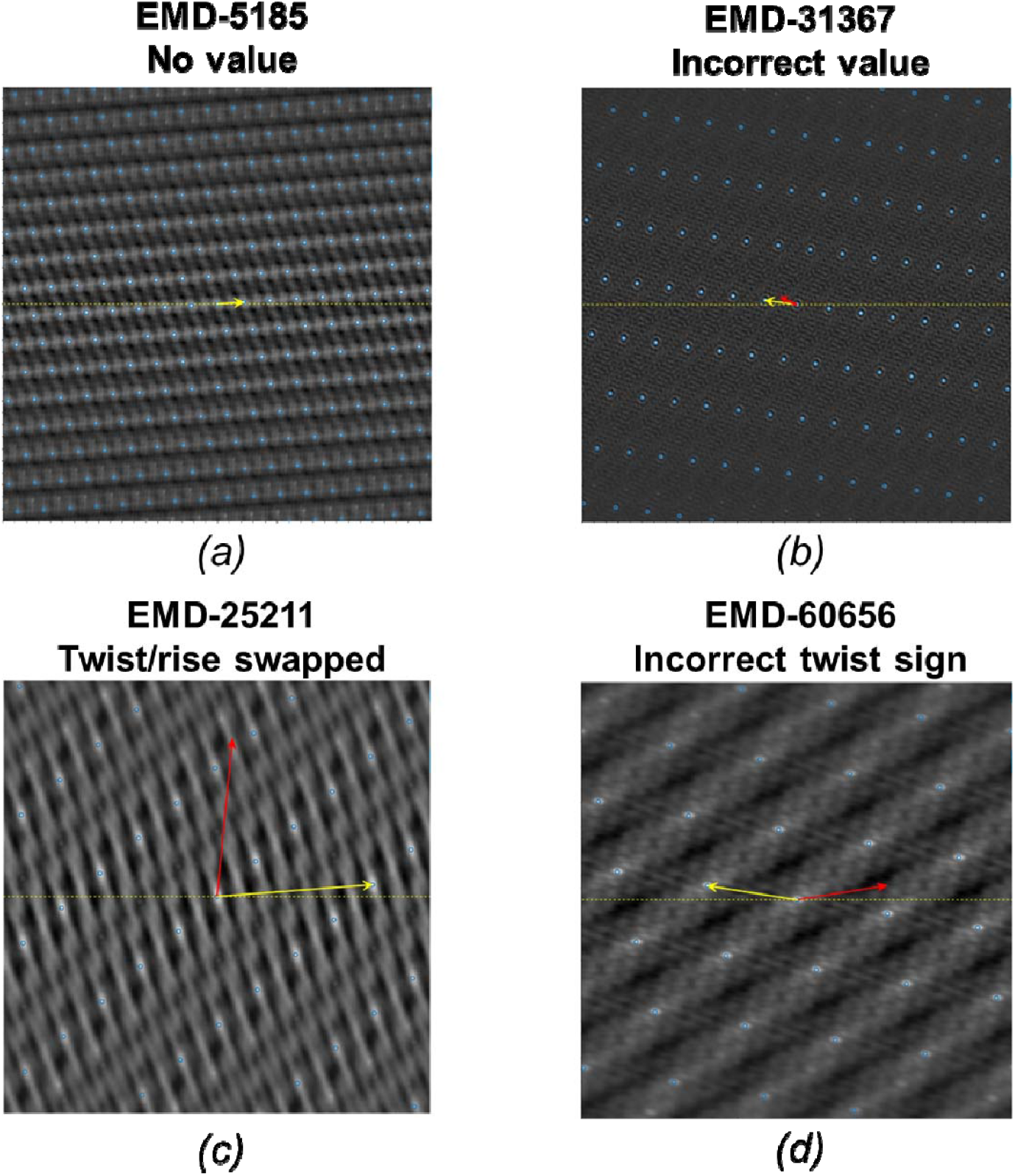
Each panel represents an example of helical reconstruction where the yellow arrow denotes the validated helical parameter determined by HI3D and the red arrow indicates the incorrect values originally deposited in the EMDB. *(a)* No values (EMD-5185): The helical parameters were missing in the original metadata. *(b)* Incorrect values (EMD-31367): The deposited helical parameters were incorrect, not related to the correct parameter, and not on a lattice point. *(c)* Twist/rise swapped (EMD-60959). *(d)* Incorrect twist sign (EMD-60656).

#### 3.2.1. Missing helical parameters (EMD-5185 (Ge & Zhou, 2011))

Some entries lack helical parameters in the metadata. An example is shown in Fig. 3A in which the yellow arrow represents the validated helical parameter. The lack of deposited helical parameters necessitates validation to fill the missing essential metadata and complete the database record.

#### 3.2.2. Incorrect values (EMD-31367 (Shan *et al*., 2021))

Some entries contain deposited helical parameters that deviate significantly from values expected from the density map. The validated value also seemed to have no direct relationship with the value deposited. In Fig. 3B, the red arrow, which represents the original deposited helical parameter, does not point to any lattice point in the HI3D lattice plot, while the validated helical parameters (marked by the yellow arrow) accurately point to the lattice point nearest to the center that corresponds to the correct twist (X-axis) and rise (Y-axis). The cross-correlation score with the validated value (0.82) is also significantly higher than the deposited one (0.04).

#### 3.2.3. Twist and rise swapped (EMD-25211 (Kreutzberger *et al*., 2022))

In multiple cases, the helical twist and rise values were ostensibly swapped. Fig. 3C illustrates what this error looks like in the HI3D lattice plot. The validated value (marked by the yellow arrow) accurately points to a lattice point, while the deposited value points to a position between the lattice points in the HI3D plot. We hypothesize that these errors were mostly from typos during deposition.

#### 3.2.4. Incorrect twist sign (EMD-60656 (Wang *et al*., 2024))

The most common error is the misassignment of the twist sign. This is a little bit tricky; it is possible that the helical parameter is correct, but the map handedness is flipped. Determining the correct handedness of a structure using cryo-EM necessitates high-resolution data and expert validation of both the atomic model and the 3D density map (Garcia Condado *et al*., 2022), which is beyond the scope of this work.

The objective of this study is limited to examining the consistency between the deposited helical parameters and the corresponding density maps. In Fig. 3D, the deposited value (red arrow) does not align with any lattice point and is on the opposite side of the validated value (yellow arrow), highlighting the importance of accurate twist sign assignment. Some of these errors might be due to typos during deposition.

#### 3.2.5. Partial symmetry (EMD-43868)

In this case, both sets of helical parameters are consistent with the deposited density map. As shown in Fig. 4A, the two arrows point to different lattice points in the HI3D plot. The yellow arrow (validated parameter) points to the lattice point (twist=65.42°, rise=5.07Å) that is closest to the equator, representing the optimal twist and rise values and the full helical symmetry. However, the red arrow (deposited parameter) points to a lattice point (twist=-0.3°, rise=111.12Å) that corresponds to a n=22 multiplier of the validated parameter. Though both helical parameters are correct, the validated helical parameter will yield more asymmetric units than the deposited helical parameter. If we could use the full helical parameter during helical reconstruction, it would potentially significantly improve the reconstruction quality and resolution.

**Figure 4.**
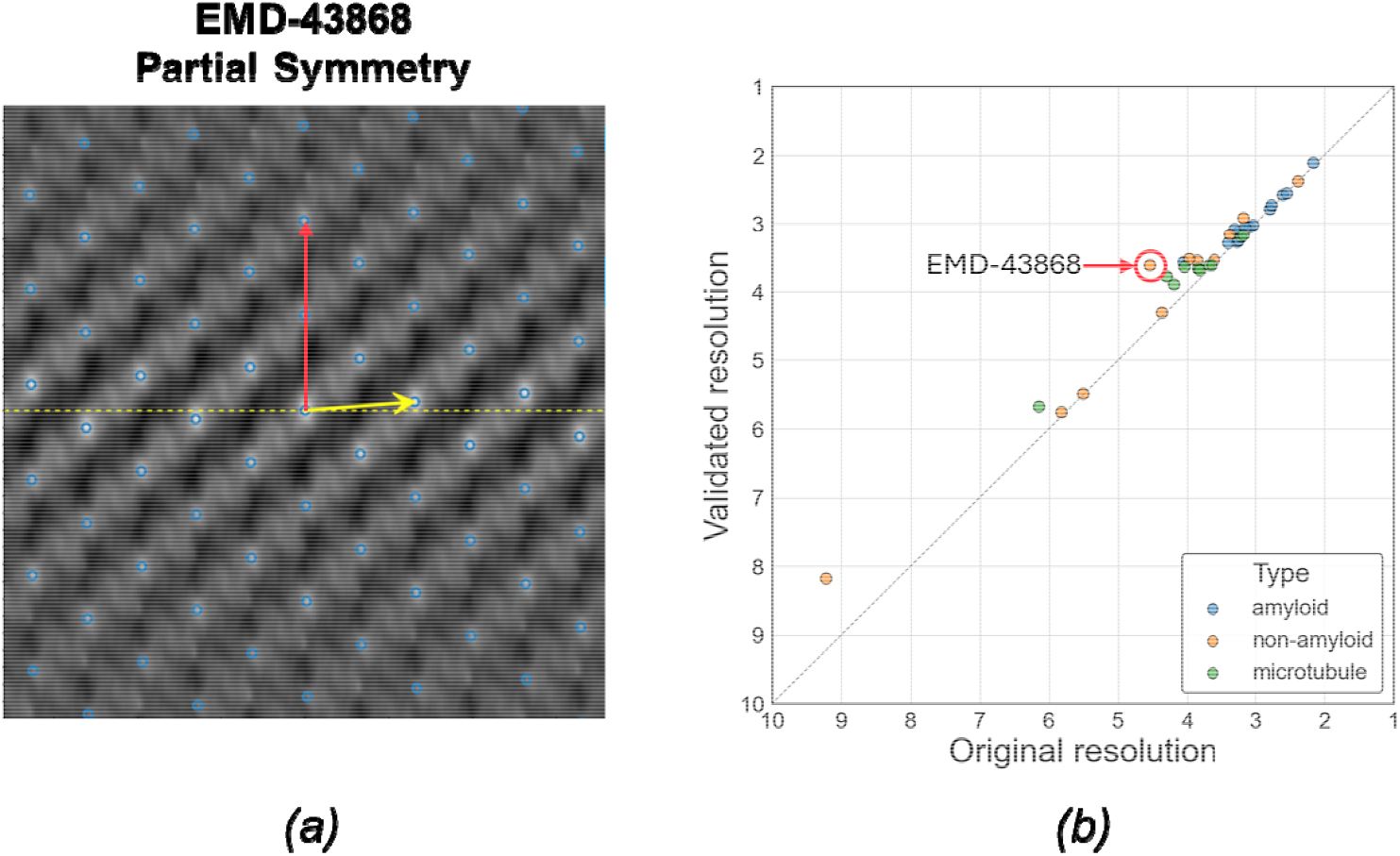
*(a)* Illustration of partial helical symmetry on a HI3D lattice plot for EMD-43868. *(b)* A paired scatter plot of map resolutions resulted from using the deposited helical parameters (X-axis) and the validated helical parameters (Y-axis). The resolution was based on the 0.143 criterion for the Fourier Shell Correlation of half maps symmetrized with a helical parameter set. Each point represents one of the 38 EMDB entries found to use partial helical symmetries in the deposited metadata.

### 3.3. Improved resolution with full helical symmetry

Our validation process identified 49 entries that potentially used partial symmetry. Since we do not have access to the original cryo-EM images used for the 3D reconstructions, we evaluated whether the map resolution could be improved by symmetrizing the deposited half-maps using the full helical symmetry identified by our validation process over symmetrization using the deposited, partial helical symmetry. As shown in Fig. 4B, the data points are consistently above the diagonal line, indicating that the resolution improved when using the validated helical parameters. Since only coherent averaging using more asymmetric units can improve map resolution, the improvements shown in Fig. 4B suggest that our validated helical parameters accurately represent the full helical symmetry.

Although most improvements are marginal or modest, there were a few entries with more significant improvements. For example, the entry EMD-43868 exhibits the greatest resolution improvement from 4.54 to 3.61 Å. We further examined the HI3D lattice plot of its density map to understand how the two sets of helical parameters are related. As shown in Fig. 4A, the deposited helical parameter points up nearly vertically and skips many lattice points closer to the equator, while our validated parameter points to the lattice point nearest to the equator and the center origin. To examine if the improvements in resolution number is supported by improved structural details, we systematically compared the density maps (Fig. 5), including the original density (Fig. 5A), the density symmetrized using the deposited helical parameters (Fig. 5B), and the density symmetrized using the validated parameters (Fig. 5C). The map symmetrized with the validated helical parameters demonstrated the best density quality (Fig. 5C).

**Figure 5.**
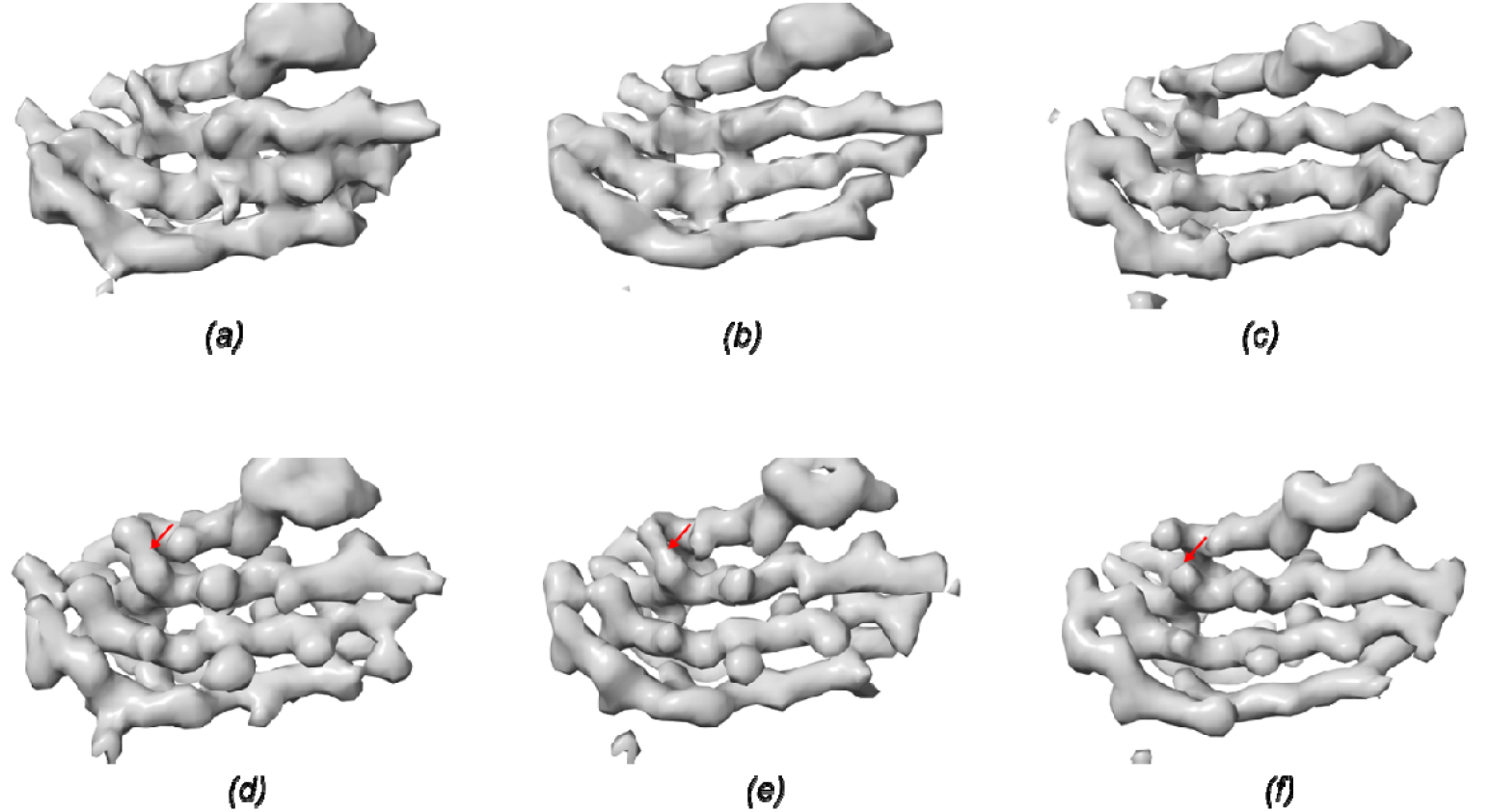
*(a)* The original density of EMD-43868. *(b)* The density in (A) symmetrized with the deposited helical parameter. *(c)* The density in *(a)* symmetrized with the full helical parameter. *(d), (e)* The density maps shown in *(a)* and *(b)*, respectively, were filtered to match the 1D structural factors profile of *(f). (f)* The density map of *(c)* auto-sharpened using Phenix. The area with clear

However, since different maps can exhibit varying voxel value ranges and the apparent structural features are influenced by subjectively chosen threshold values (Beckers *et al*., 2019), we further ruled out these potential factors to objectively and fairly compare these three maps. We first used Phenix (Afonine *et al*., 2018) to sharpen the density symmetrized with the validated parameters, ensuring optimal visual representation (Fig. 5F). The other two maps were then filtered to match the 1D structural factor radial profile of the sharpened map (Fig. 5D,E). With these steps, the maps were brought to the same filtering level to allow visualization using the same threshold, and the visual differences would be from true structural differences. It could be seen that the map symmetrized using the validated helical parameters (Fig. 5F) retained the best quality, indicated by a cleaner beta-strand separation. The example shown here (Fig. 4A and Fig. 5) is to illustrate the significance of utilizing the full symmetry. To truly verify the full helical symmetry and to maximize the resolution and map quality, access to the original micrographs and reprocessing the data with helical reconstruction using the full symmetry will be needed.

## 4. Discussion

In this study, we developed a systematic validation strategy for the helical symmetry parameters, used this strategy to validate the helical parameters of all EMDB helical structure entries (2,025 in total, as of April 25, 2025), and identified a considerable number of inconsistencies between the deposited 3D density maps and their associated helical parameters. These inconsistencies fall into multiple categories, including missing helical parameters, swapped twist and rise values, incorrect twist signs, and bona fide errors. These errors were corrected using a rigorous validation process. Partial symmetries, which are correct but suboptimal, were also identified for some entries. Further symmetrization with the full helical symmetry was shown to improve the resolution, although only modestly for most entries, whilst raw cryo-EM images were not available. An example with most improvement from 4.54 to 3.61 Å and noticeably better resolved structural features highlights the importance of using the full helical symmetry, particularly during 3D structural determination from 2D images.

In summary, the systematic validation process described here could identify and correct helical parameter errors in the EMDB, ensuring consistency between the helical parameters and the corresponding 3D density maps. The availability of validated helical parameters will be valuable for downstream applications, such as training deep learning models on helical structures within EMDB. The validation protocol will be valuable for individual investigators to ensure the usage of correct and full helical symmetry to achieve the best resolution and map quality, and for EMDB to consider adopting during the curation process post-deposition of a helical structure to ensure the correctness of the helical symmetry parameters and avoid further propagations of potential errors.

## Acknowledgements

This work was supported in part by NIH grants RF1NS110437, R01AG071177, R21AG081686, Purdue Showalter Scholars award, and Penn State startup fund (WJ). We thank the Jiang lab members’ help during the validation process.

## Appendix A.

**The complete list of validated helical parameters for the 2**,**025 helical structures in EMDB as of April 25, 2025**

In the HI3D_link column, clickable links are provided to show the HI3D lattice plot with both the original deposited helical parameters (red arrow) and the validated helical parameters (yellow arrow). If the arrow does not point to a lattice point, the corresponding helical parameters are incorrect. If the arrow points to a lattice point but not the one closest to the equator line and the center origin, the corresponding helical parameters are correct but represent partial helical symmetries.

